# Non-canonical role of Hippo tumor suppressor serine/threonine kinase 3 STK3 in prostate cancer

**DOI:** 10.1101/2021.02.21.432137

**Authors:** Amelia U. Schirmer, Lucy M. Driver, Megan T. Zhao, Carrow I. Wells, Julie E. Pickett, Sean N. O’Bryne, Benjamin J. Eduful, Xuan Yang, Lauren Howard, Sungyong You, Gayathri R. Devi, John DiGiovanni, Stephen F. Freedland, Jen-Tsan Chi, David H. Drewry, Everardo Macias

## Abstract

Serine/threonine <u>k</u>inase <u>3</u> (STK3) is an essential member of the highly conserved Hippo Tumor suppressor pathway which regulates Yes 1 Associated protein (YAP1) and TAZ. STK3 and its paralog STK4 initiate a phosphorylation cascade that regulate YAP1/TAZ activation and degradation, which is important for regulated cell growth and organ size. Deregulation of this pathway leads to hyper-activation of YAP1 in various cancers. Counter to the canonical tumor suppression role of STK3, we report that in the context of prostate cancer (PC), STK3 has a pro-tumorigenic role. Our investigation started with the observation that STK3, but not STK4, is frequently amplified in PC. A high STK3 expression is associated with decreased overall survival and positively correlates with androgen receptor (AR) activity in metastatic castrate resistant PC. XMU-MP-1, an STK3/4 inhibitor, slowed cell proliferation, spheroid growth and matrigel invasion in multiple models. Genetic depletion of STK3 decreased proliferation in several PC cell lines. In a syngeneic allograft model, STK3 loss slowed tumor growth kinetics *in vivo* and biochemical analysis suggest a mitotic growth arrest phenotype. To further probe the role of STK3 in PC, we identified and validated a new set of selective STK3 inhibitors, with enhanced kinase selectivity relative XMU-MP-1, that inhibited tumor spheroid growth and invasion. Consistent with the canonical role, inhibition of STK3 induced cardiomyocyte growth and had chemo-protective effects. Our results contend that STK3 has a non-canonical role in PC progression and inhibition of STK3 may have therapeutic potential for PC that merits further investigation.

**Significance:** Our findings illuminate a new actionable target for PC therapy that would traditionally be overlooked due to its canonical role as a tumor suppressor in other cancer types.

## Introduction

Prostate cancer (PC) remains the second leading cause of cancer related deaths in men. Metastatic castration resistant PC (mCRPC) occurs when PC progresses despite continuous androgen deprivation therapy. This is largely due to aberrations in the Androgen Receptor (AR), including AR gene amplification, expression of splice variants, or mutations. Over the past few years, a number of AR targeted therapies have been FDA approved, including enzalutamide and abiraterone acetate for mCRPC, and most recently darolutamide and apalutamide. However, in spite of these life prolonging therapeutic advances, there is no cure for mCRPC, resulting in patients eventually succumbing to the disease. Thus, there is a need to identify other molecular targets for mCRPC beyond AR.

We have focused our efforts on understudied kinases that are correlated with poor cancer patient outcomes in PC. Kinases are high value targets due to their druggability and have emerged as the most successful drug target of the 21st century, with more than 60 small molecule kinase inhibitors approved by the FDA for oncology (1). This enzyme family contains more than 500 members, presenting a wealth of opportunities for the continued introduction of life extending medicines. Yet, a large number of kinases are understudied and thus overlooked as potential therapeutic targets (2,3). Herein we investigate the role of Serine/Threonine Kinase 3 (STK3) also known as STE20-Like Kinase MST2 a key member of the Hippo Tumor Suppressor signaling pathway.

The term “Hippo” is derived from the overgrowth phenotype in drosophila that is associated with genetic deletion of the *hpo* gene which encodes a Ste-20 family protein kinase that restricts cell proliferation and regulates apoptosis (4). The mammalian homolog *STK3* gene and its paralog *STK4*, encode what are commonly referred to as MST2 and MST1 kinases, respectively. STK3 and STK4 are essential members of the highly conserved Hippo Tumor Suppressor pathway (4,5) (*hereafter, simply STK3 and STK4 kinases*). Mammalian STK3/4 kinases bind to adaptor protein Salvador homolog 1 (SAV1), phosphorylate large tumor suppressor homolog 1/2 (LATS1/2) and Mob kinase activator (MOB1). Phosphorylation of homologous transcriptional co-activators <u>Y</u>es <u>A</u>ssociated <u>p</u>rotein (YAP) and transcriptional co-<u>a</u>ctivator with PD<u>Z</u> binding motif (TAZ) by LATS1/2 ensues which leads to their cytoplasmic retention or degradation (6). Inactivation or loss of Hippo kinase signaling leads to non-phosphorylated active YAP/TAZ, which promotes cell proliferation and inhibits apoptosis. YAP/TAZ transcriptional oncogenes are hyperactivated in many cancer types (6,7). Hyper-activation of YAP in PC has been observed in a number of studies (8). Thus, the consensus would be that loss of STK3/4 Hippo kinase signaling results in hyperactivation of YAP/TAZ. However, our analysis of genomic data shows that STK3, but not STK4, is amplified human PCs and correlates with worse patient outcomes. Thus, we hypothesized that STK3 may have a non-canonical pro-tumorigenic role in PC.

Given the high value of STK3 as an actionable target and the urgent need for new molecular targets in PC, we sought to investigate a potential non-canonical pro-tumorigenic role of STK3. To complement STK3 molecular genetic studies, we utilized available STK3/4 inhibitor XMU-MP-1 and identified novel complementary series of STK3 small molecule inhibitors with different structures, narrower kinase inhibition profiles, and more potent cellular activity. These small molecules may also be useful tools for the research community to elucidate STK3 function in other disease contexts. Our cumulative data support that STK3 has a non-canonical pro-tumorigenic role in PC, which may be targeted by small molecule inhibitors to reduce PC growth and progression.

## Materials and Methods

### Cell lines and culture conditions

<u>H</u>i-<u>M</u>yc <u>v</u>entral <u>p</u>rostate <u>2</u> cells (HMVP2) were a gift from Dr. John DiGiovanni (9). C4-2 cells were obtained from the Chen Lab at Duke University. Cell lines were authenticated by STR profiling and tested for mycoplasma at Duke University cell culture facility by MycoAlert PLUS test Lonza. DU145 and 293T cells were grown in Dulbecco’s Modified Eagle Medium (DMEM) + 10% fetal bovine serum (FBS). PC3, 22RV1, HMVP2, C4-2 and LNCaP cells were cultured in RPMI + 10% FBS. LAPC-4 cells, a gift from Dr. William Aronson at UCLA, were cultured in Iscove’s Modified Dulbecco’s Medium (IMDM) + 10% fetal bovine serum (FBS) + 1nM R1881. For androgen deprivation charcoal stripped FBS (CSS)(MilliporeSigma, St. Louis, MO., USA) was used instead of FBS in media.

### Western blots

Cells were collected in radioimmunoprecipitation assay buffer (RIPA) lysis buffer with protease cocktail inhibitor 1 and phosphatase cocktail inhibitor 2 and 3 (MilliporeSigma). For tumor biochemical analysis tumoral tissues were ground in liquid nitrogen with a mortar and pestle and then extracted with RIPA. Protein was quantified using DC Bio-Rad protein assay (Bio-Rad, Hercules, CA., USA) and 30 to 60 μgs of protein were run on SDS-PAGE gels and transferred onto 0.45 or 0.22 μmol/L nitrocellulose. Antibodies p-MOB1(Thr35), MOB1, Cyclin D1, p-YAP(Ser397), p-YAP(Ser127), YAP, STK4, TAZ, CDK1(Tyr15), p-STAT3 (Cell Signaling Technology, Danvers, MA USA) were all used at 1:1,000 dilution. Antibodies for GAPDH and AR (Cell Signaling Technology) used at 1:4,000 and 1:2,000 dilution respectively. Antibody for STK3 (ThermoFisher, Waltham, MA., USA) used at 1:2,000 dilution. For Western blot quantification, ImageJ 1.52a software was utilized to quantify band densitometry and Actin or GAPDH loading controls for normalization. When shown as percentage, treatment group ratios were divided by control group ratio (100%) and multiplied by 100.

#### Cell proliferation assays

For two-dimensional proliferation assays, cells were plated in a 96-well clear walled, clear bottom plates and after 24 hours, cells were treated as specified for 120-192 hours using DMSO as a vehicle. For three-dimensional spheroid assays, cells were plated in 96 well, clear walled, ultra-low attachment plates and after 24 hours when spheres formed, cells were treated for 144-192 hours with DMSO as a control. Proliferation studies were conducted on an IncuCyte S3 live-cell imaging system (Sartorious, Ann Arbor, MI., USA) on 96 whole-well scan mode. Plates were imaged every 8-24 hours and quantified with bundled image analysis software. For real-time cell death assays, Incucyte® Cytotox Green Reagent (Satorius) was used to indicate dead cells.

### Gene Knockdown

For inducible knockdown of STK3 shERWOOD UltramiR lentiviral Inducible shRNA system was used (Transomics Technologies Inc., Huntsville, AL., USA). To deplete mouse STK3 expression in HMVP2 cells TRC2 MISSION shRNAs in pLKO.1-puro vector backbone were used (Millipore-Sigma). Lentivirus was generated with psPAX2 and pMD2.G packaging plasmids using with the Polyplus jetPRIME transfection kit in 293T cells. Virus was collected at 48 hours post transfection and a 1:1 ratio of viral media to fresh culture media with 8ug/mL of polybrene infection reagent was incubated for 24 hours on target cells.

### Kinetic cell migration and matrigel invasion

Scratch wound migration and matrigel invasion assays were conducted with an IncuCyte S3 live-cell imaging system (Sartorius Bioscience). DU145 cells (4.5 × 10^5^) were plated on imagelock 96-well plates and treated for 24 hours with indicated drug. Cells were then washed twice with PBS, and a 96-pin wound making tool was used to make a uniform scratch in all 96 wells simultaneously (IncuCyte 96-well WoundMaker Tool, cat. # 4563). For cell migration (no matrigel) assays, complete DMEM with or with out treatment was added to wound with indicated treatments and plate imaged over time. For matrigel invasion assays, scratch wound was overlaid with 3 mg/mL of matrigel and incubated for 1 hour at 37°C, and then overlaid with 100 μL of complete DMEM with indicated treatments and imaged over time. Quantification was conducted using the IncuCyte Scratch Wound software module. For three-dimensional spheroid invasions, 500 cells were plated in PrimeSurface 3D culture, ultra-low attachment, 96 well, U bottom, clear plates 50 μL of media for 24 hours to allow spheres to form (S-BIO, Hudson, NH USA). Then matrigel was gently mixed for final concentration of 3 μg/mL and incubated for 1 hour at 37°C, and then overlaid with 100 μL of RPMI media and indicated treatment. Quantification was conducted using the IncuCyte spheroid software module.

### *In vivo* syngeneic allograft study

Mouse experiments were conducted as approved by the Duke University IACUC board protocol number IACUC006565. Briefly, 32 male FVB/N mice aged 6 weeks (Charles River Laboratories Raleigh NC, USA) were injected with HMVP2 spheroids grown in ultralow attachment plates as previously described (9). Approximately 2×10^5^ cells were injected in 1:1 RPMI to matrigel per animal subcutaneously in the right flank. Caliper measurements were commenced 14 days after inoculation three times per week and animal body weights measured twice a week. Tumor volumes were calculated using the formula L x (W)^2^/2. At terminal endpoint, wet tumor weights were recorded, tumoral tissues divided and fixed in formalin or flash froze in liquid nitrogen.

### NanoBRET Assays

Intracellular NanoBRET assays were performed as previously described (10) using NanoLuc®-STK3 Fusion Vector Cat.# NV4301 (Promega, Madison, WI, USA). Briefly, NanoLuc-STK3 along with carrier DNA was transfected into HEK-293 cells in a 96 well format. NanoBRET Tracer K10 was used at a concentration of 1 μM to test compounds at 11 concentrations. To measure NanoBRET signal, NanoGlo substrate (Promega) at 1:166 to Opti-MEM media in combination with extracellular NanoLuc Inhibitor (Promega) diluted 1:500 was combined to create a 3X stock solution. A total of 50 μL of the 3× substrate/extracellular NanoLuc inhibitor was added to each well. The plates were read within 10 minutes on a GloMax Discover luminometer (Promega). Biological replicates were normalized and fit using a sigmoidal, 4 parameter logistic regression binding curve in Prism Software (GraphPad, La Jolla, CA, USA). IC_50_ and standard error (SE) were calculated in Prism Software.

### Enzyme Assays

Assays were performed at Eurofins discovery services (Celle-L’Evescault, France) as previously described (11). Investigational compounds were provided as 10 mM stock solutions and screened in 10-point dose–response at the Km of ATP using 2-end recombinant STK3 protein and myelin basic protein as a substrate.

### Compound Kinase Selectivity

KINOMEscan scanMax assay panel was performed at Eurofins DiscoverX (San Diego, CA., USA) across 403 wild-type human kinases and 65 mutant human kinases at a compound concentration of 1 μM as previously described (12). Selectivity score 10 or S_10_ (1 µM) was used for compound selectivity metric, where the threshold is 10% of activity remaining relative to control for a given kinase. S-score is calculated by dividing the number of inhibited protein kinases at or above this threshold, by the total number of tested protein kinases.

## Statistical analysis

GraphPad Prism 9.0.0 was used for statistical analysis of *in vitro* studies. For time-course live cell proliferation assays and migration/invasion assays, 2-way repeated measures ANOVA multiple comparison analysis was conducted with Dunnett’s multiple comparison test for adjusted p-value both with alpha=0.05. Tumor growth kinetics *in vivo* were analyzed using a generalized estimating equation with exchangeable correlation to test if tumor volume growth varied over time between the three treatment arms. Time was treated as a categorical variable to not assume linear growth. P<0.05 was considered statistically significant. Copy number alteration (CNA), transcriptome profiles, and clinical information of patient data were downloaded from cBioPortal and Log-rank Mantel-Cox test was performed to estimate significance of survival difference and hazard ratio between stratified groups. For gene correlation studies we used a web-based software named the Prostate Cancer Transcriptome Atlas (PCTA)(13). Androgen_Response hallmark gene set activation score was computed by using Z-score method (14). Correlations were computed by Spearman’s rank correlation method.

## Results

### STK3 gene is amplified in PC

Analysis of genomic data from 158 non-redundant studies in the cBioPortal with a cut off of ≥10% copy number alteration (CNA) frequency shows that STK3 is amplified in a number of cancer cohorts (Fig. 1A) (15). Of the 14 cohorts with ≥10% STK3 amplification, 4 were PC cohorts. In contrast, no PC cohort showed a ≥10% amplification of STK4 (Fig 1B).

**Figure 1.**
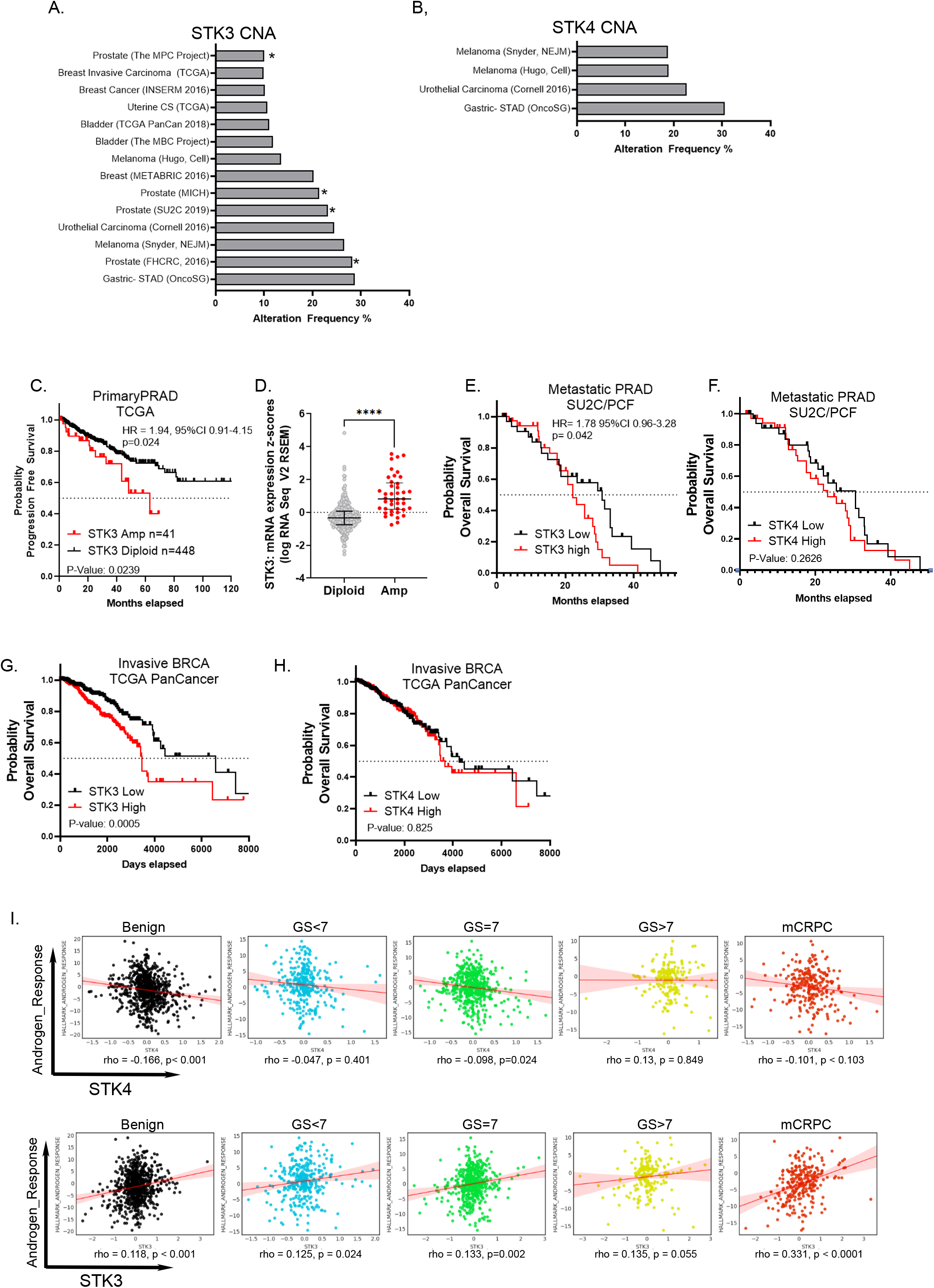
Correlative analysis of STK3 and PC patient outcomes. A) Cancer cohorts with >10% STK3 and B) STK4 copy number amplification. * denotes PC cohorts. C) Progression free survival analysis of STK3 diploid and STK3 amplified patients from TCGA/PRAD, p=0.0239. D) STK3 mRNA express z-scores in diploid and STK3 amplified PCs. ****p<0.0001. E) Survival analysis of advance mCRPC by lower and upper median STK3 expression and F) low and high STK4 expression groups. Survival analysis of BC patient overall survival in patients stratified by upper (n=503) and lower (n=503) median expression z-scores of G) STK3 and H) STK4. I) Spearman’s rank correlative analysis of STK4 and STK3 and AR response gene signature. GS=Gleason Score, mCRPC=metastatic castration resistant PC. Rho and p-values denoted on graph.

### STK3 correlates with worse outcomes in PC

Progression free survival analysis of patients from the TCGA prostate adenocarcinoma cohort (n=489) shows significant difference in time to progression in STK3 amplified patients (n=41) compared to STK3 diploid patients (n=448) (HR = 1.94, 95%CI 0.91-4.15, p=0.024, Log-rank) (Fig. 1C). TCGA prostate adenocarcinomas with increased STK3 gene copy number were associated with increased STK3 mRNA levels (Fig. 1D). Next, we queried patients with available RNA-Seq gene expression and survival data from the metastatic Stand Up 2 Cancer/Prostate Cancer Foundation cohort (SU2C/PCF) (16). Patients were stratified into lower median (STK3 low, n=36) and upper median STK3 expression (STK3 high, n=35) groups by mRNA expression z-scores for survival analysis. Compared to STK3 low expression group, STK3 upper median expression group had a significantly decreased survival rate, median survival 30.7 vs 22.2 months, respectively (HR= 1.78, 95%CI 0.961 to 3.218, p= 0.042, Log-rank) (Fig. 1E). Importantly, when we stratified patients by STK4 median expression, there was no statistically significant difference in survival between groups (Log-rank p= 0.263) (Fig. 1F).

Given STK3 is frequently amplified in breast cancer (BC), we also queried BC outcomes (Fig. 1 G-H). Patient samples from the TCGA Invasive Breast Cancer cohort stratified by STK3 above the median upper expression levels compared to lower STK3 expression showed significantly decreased overall survival (HR= 1.84, 95%CI 1.314 to 2.580 p= 0.0005, Log-rank). When BC patients were stratified by STK4 median expression, there was no significant difference in overall survival (Log-rank p=0.825).

### STK3 gene expression is correlated with AR activity in mCRPC

To determine if STK3 plays a role in AR dependent or independent PCs, we next ran correlative analysis of STK3 or STK4 gene expression and AR response signature (HALLMARK_ANDROGEN_RESPONSE) across PC disease states. For STK4 and AR response, there was a negative or no significant correlation observed (Fig. 1I). However, STK3 gene expression and AR response gene signature had a significantly positive correlation at every disease state with the highest correlation in the mCRPC setting (Spearman’s rho value =0.33, p= 0.001). Altogether, analysis of STK3 in human PC specimens suggest that STK3 may play a role in lethal AR driven mCRPC.

### STK3 expression in PC cell lines

Across commonly used PC cell lines, there was a trend for an inverse relationship between STK3 and STK4 protein expression (Fig 2A). In addition, in AR positive cell lines STK3 tends to be higher and YAP/TAZ expression lower. Exploring PC cell lines for syngeneic studies, we found Hi-Myc Ventral Prostate 2 (HVMP2) cell line derived from the commonly Hi-Myc transgenic PC mouse model, has significantly elevated levels of STK3 (Fig. 2A).

**Figure 2.**
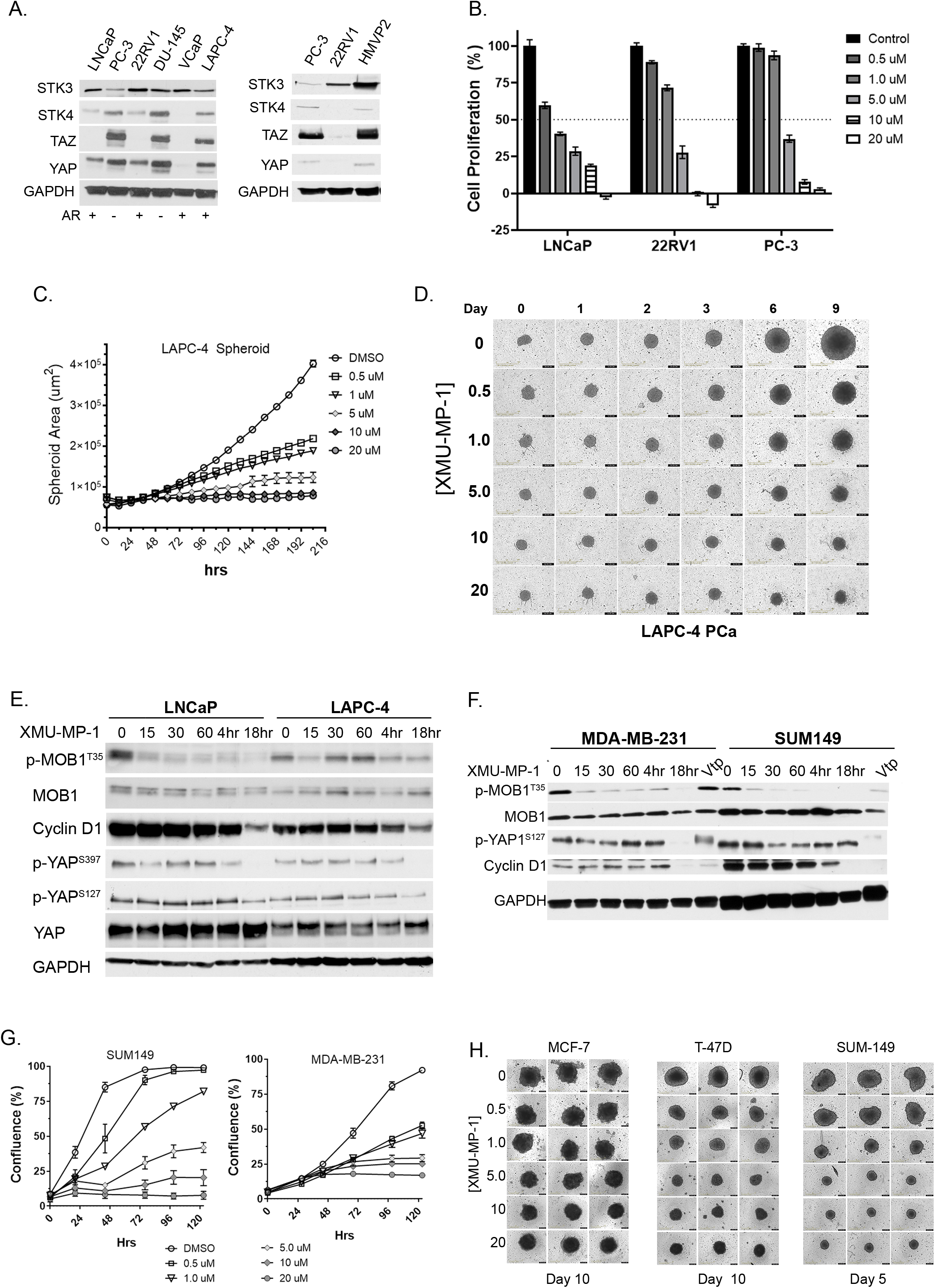
Pharmacological inhibition of Hippo Kinases in PC and BC. A) Expression of STK3 and STK4 across PC cell lines. B) Cell proliferation assessed by total cell confluence normalized to vehicle control of 22RV1, PC-3 and LNCaP cells treated with XMU-MP-1 (n=5/dose), data shown 96hrs post treatment. C) Graphical representation of LAPC-4 spheroid growth kinetics (spheroid area) over time and D) representative bright field images of LAPC-4 spheroids treated as denoted with XMU-MP-1 (n=5). E) Western blot analysis of denoted PC and F) BC cell lysates treated with 10 µM XMU-MP-1 collected at indicated time-points post treatment. Vtp = verteporfin. G) Assesement of BC cell proliferation and H) BC spheroid growth of three cell lines with various doses of XMU-MP-1 (n=5/dose).

### STK3/4 inhibitor, XMU-MP-1, slows PC and BC cell growth and matrigel invasion

First, we wanted to determine if STK3 is a potential druggable target in PC. To this end, we utilized the available STK3/4 small molecule inhibitor XMU-MP-1 across a battery of cell lines (17). In LNCaP, 22RV1 and PC-3 cells, we observed a dose dependent decrease in proliferation rates with XMU-MP-1 treatment (Fig. 2B). In addition, growth of LAPC-4 3D spheroids was significantly blunted with XMU-MP-1 in a dose dependent manner (Fig. 2C-D). Western blot analysis of LNCaP and LAPC-4 cell lysates treated with 10 μM XMU-MP-1 showed decreased levels of p-MOB1 and phospho-YAP at both Ser127 and Ser397, denoting activated YAP, yet PC cells were growth inhibited (Fig. 1). Consistent with slowed proliferation rates, cell cycle progression marker cyclin D1 was down regulated 18hrs post XMU-MP-1 treatment.

Given that STK3 is also frequently amplified in BC (Fig 1A), we also tested the effects of XMU-MP-1 on BC cells. Similar to our observations in PC cell lines, western blot analysis of BC cell lines treated with XMU-MP-1 also displayed reduced phosphorylation of YAP, but reduced cell proliferation marker Cyclin D1 (Fig. 2F). Verteporfin, a know YAP inhibitor treatment for 18hrs was used as a positive control. In longitudinal proliferation assay, MDA-MB-231 and SUM-149 BC cells, were also growth inhibited in a dose dependent manner (Fig. 2G). Similar results were observed in various three-dimensional BC spheroid models (Fig 2H). Together, data from PC and BC cells, shows that STK3/4 inhibition with XMU-MP-1 results in reduced cell proliferation despite YAP1 activation (reduced phosphorylation).

### Genetic depletion of STK3 slows PC cell proliferation

We utilized shRNA to test if STK3 is essential for PC cell proliferation (Fig 3A and SX). Western blot validation of STK3 inducible knockdown coincides with reduced levels STK3 phospho-target p-MOB1 Thr35. Importantly, STK4 levels are not induced to compensate for STK3 loss. In addition, we observed increased levels of cell cycle inhibitor p27. Modest increases in YAP1 Ser127 were observed but appear in non-targeting doxycycline treated cells as well. This indicates that YAP is not activated in this model due to loss of STK3. In LNCaP, C4-2 and HMVP2 cell lines, STK3 knockdown with two shRNAs compared to non-targeting shRNA control, resulted in significantly reduced proliferation rates (Fig. 2B-C). In LAPC-4 3D spheroid model, STK3 gene knockdown also resulted in slowed tumor spheroid growth kinetics (Fig. 3D-E).

**Figure 3.**
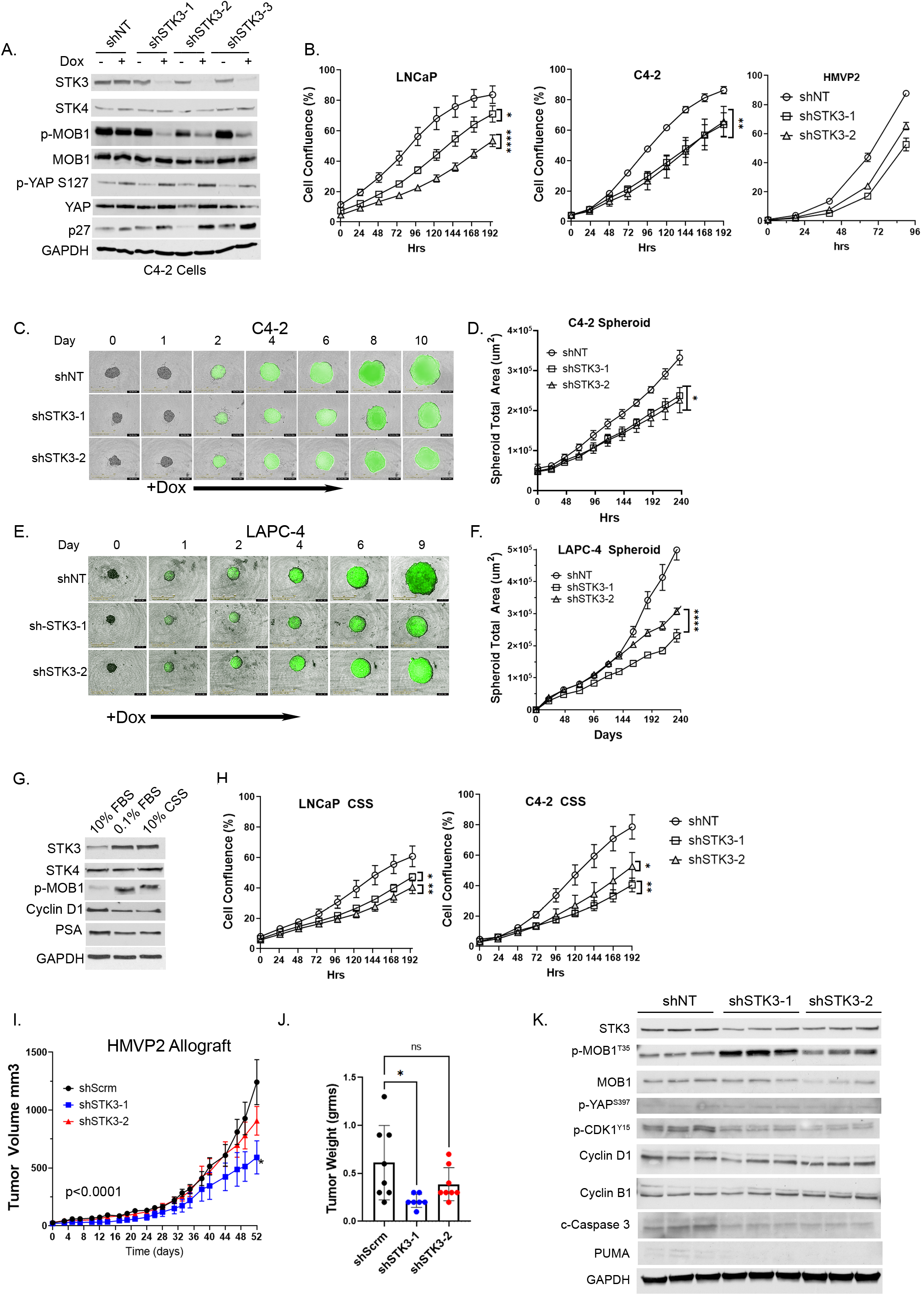
Assessment of STK3 genetic depletion in PC cells. A) Western blot validation of STK3 shRNA inducible knockdown (doxycycline 0.5 ug/ul) and assessment of denoted proteins. B) Cell proliferation assays of LNCaP, C4-2B and C) HMVP2 PC cells with shNT or shSTK3s (n=4). D) Time course spheroid growth in LAPC-4 with shNT or shRNAs for STK3. E) Western blot analysis of LNCaP cell lysates grown in varying serum conditions for 72hrs. F) Cell proliferation assays of LNCaP and C4-2B grown in CSS (n=5). G) Phase images of C4-2B cells with indicated shRNA at 192hrs. I) Longitudinal growth kinetics of HMPV2 allograft tumors with shNT (n=8), shSTK3-1 (n=8), shSTK3-2 (n=7), interaction time vs shRNA arm p<0.001. Pairwise comparison shSTK3-1 vs. shNT: p<0.001; shSTK3-2 vs. shNT: p=0.23. J) Final tumor wet weight. K) Western blot analysis of pooled tumor lysates (n=6/shRNA) loaded in technical triplicate.

### STK3 in castrate growth conditions

Given that STK3 is amplified in advanced mCRPC cohorts, we asked whether STK3 plays a role in hormone independent PC cell growth. For these studies we utilized charcoal stripped serum (CSS) to mimic androgen deprivation conditions and as a control we used low serum growth conditions. Interestingly, after 72hrs of growth in CSS, STK3 protein levels and activity (p-MOB1) were increased (Fig. 3F). Consistent with nutrient/hormone deprived growth conditions we found reduced levels of cell cycle marker cyclin D1 and AR target gene PSA. In both LNCaP and C4-2 cell lines grown in CSS, STK3 loss resulted in blunted proliferation compared to non-targeting control (Fig. 3G). Morphologically, C4-2 cells with STK3 knockdown, when compared to shNT cells, had increased cytoplasmic volume and were multinucleated (Fig. 3H). Together, these data indicate that STK3 plays a role in castration resistant PC cell proliferation. Thus, STK3 may impart growth advantage under androgen-deprived growth conditions.

### STK3 knockdown slows allograft tumor growth *in vivo*

To determine the role of STK3 on PC growth *in vivo* we utilized HMVP2 syngeneic allograft model. Loss of STK3 *in vivo* showed an overall statistically significant interaction (p<0.0001) between treatment arm and time, which tests whether tumor volume differs by treatment (i.e. shSTK3 vs shNT) over time (Fig. 3I). On pairwise comparison, shSTK3-1 (p<0.001), but not shSTK3-2 (p=0.23) was significantly different compared to shNT control arm. At the experimental endpoint, similar results were observed with tumor weights (Fig. 3J). Biochemical analysis of pooled tumor protein lysates (n=6 tumors per group), confirmed reduced levels of STK3 in shSTK3 tumors compared to shNT control tumors (Fig. 3K). Interestingly, we did not observe reduced levels of p-MOB1 in STK3 depleted tumors. However, loss of STK3 *in vivo* resulted in reduced levels of p-CDK1 Tyr15, a previously described STK3 downstream target (18). In addition, we observed decreased levels of cleaved Caspase 3 and the proapoptotic protein PUMA in shSTK3 tumors compared to shNT tumors. These data suggest that STK3 loss *in vivo* results in slowed tumor growth kinetics due to cell cycle alterations and not cell death due to hyper-activation of YAP as described in other models (19).

### Identification and validation of novel STK3 small molecule inhibitors

To identify new medicinal chemistry starting points for targeting STK3, our team screens diverse kinase inhibitors in a large assay panel, and carefully scruitinized previously published kinome data (20). Through this literature mining, we identified two independent scaffolds (Fig. 4A) disclosed as inhibitors of well-studied kinases that also possess what we perceived to be exploitable off-target STK3 inhibition (21,22). One lead, alias UNC-BE4-017, referred to as compound A1 hereafter, is a pyrrolopyrazine that was originally disclosed in a medicinal chemistry campaign targeted at the JAK kinases (22). The compound potently inhibits JAK1, 2, and 3, but only 3 kinases, one being STK3, out of the panel of 48 are inhibited >90% indicating good kinase selectivity for a starting point. The second lead, alias UNC-SOB-5-16, referred to as compound B1 hereafter, a pyrrolopyrimidine, came from a literature series designed to target the kinase LRRK2 (21). It is also an inhibitor of STK3, with an IC_50_ = 22 nM and in the DiscoveRx KINOMEscan panel of 451 kinase assays only 10 kinases give a percent of control (PoC) <90 at 1µM. Thus, compounds A1 and B1 are very selective across the kinome with only a handful of off-targets. We synthesized these lead compounds and some additional analogs from each series to test for STK3 engagement and inhibition (**Supplemental Materials and Methods**).

**Figure 4.**
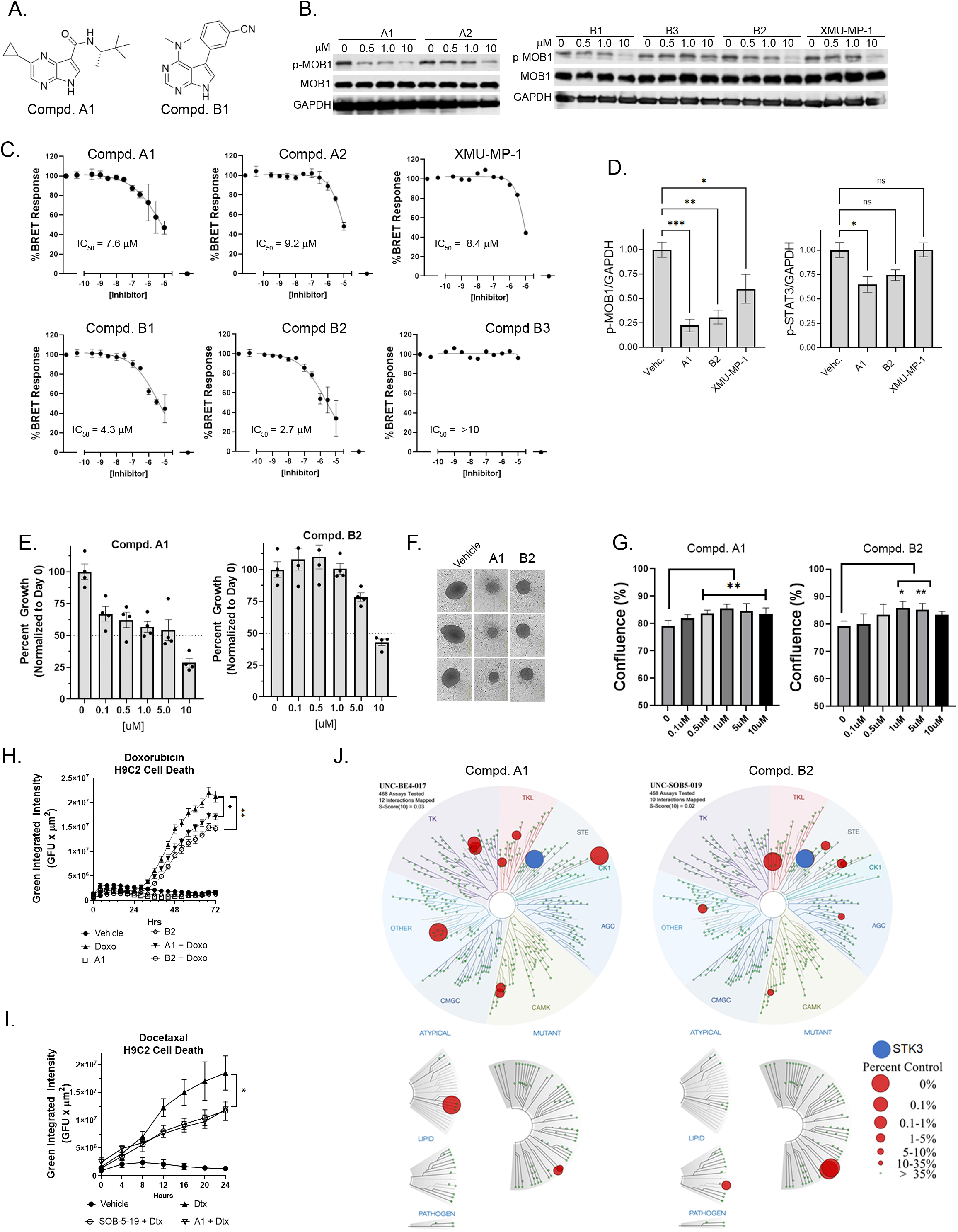
Identification and assessment of STK3 small molecule inhibitors. A) Structure of lead compound A1 and B2. B) Western blot analysis of STK3 target p-MOB1 in HMVP2 cells treated with denoted compound for 30 minutes. C) STK3-NanoBRET intracellular 11-point IC50 dose response curves with denoted compound. D) p-MOB1 and p-STAT3 ratios normalized go GAPDH relative to vehicle control (n=4). ns= non-significant, *p=0.0473, **p=0.0017, ***p<0.0007. E) Efficacy of lead STK3 compounds on HMVP2 spheroid growth at Day 9 (n=4). F) Representative bright field images of vehicle or 10 μM treated HMVP2 spheroids at 9 day endpoint. G) H9C2 cardiomyocyte proliferation assay treated with denoted compounds 72hrs post treatment (n=4) *p<0.05, **p< 0.01. H) H9C2 proliferation and cell death monitored by cytotox green dye (GFP) over time treated with vehicle, doxorubicin (Doxo, 500 nM) or STK3 inhibitors (1 μM). I) Kinome tree for A1 and B2 at 1 μM doses. Kinases with 0-35% percent of control remaining activity depicted. STK3 denoted in blue.

### Validation of lead scaffolds and STK3 target engagement

First we utilized p-MOB1 Thr35, a widely validated STK3/4 phospho-substrate as a readout for STK3 inhibition in cells (23). As shown in Fig. 4B, 30 min treatment with compounds A1, A2, B1, B2 and XMU-MP-1 resulted in reduced STK3 activity at the 10 μM dose. However, A1 and B2 were more potent, blocking MOB1 phosphorylation at lower doses (Fig. 4B). We next employed an STK3-NanoBRET assay to quantitate STK3 occupancy and affinity of our compounds in live cells (10). In STK3-NanoBRET assay, A1 and B2 were found to have the lowest in cell IC_50_ values, 7.6 and 2.7 μM, respectively (Fig. 4C). XMU-MP-1 was found to have an IC_50_ of 8.4 μM, which coincided well with western blot results. Of note, B3 showed no activity in western blot assays and an IC_50_ out of range (>10 μM) in the STK3 NanoBRET assay, indicating strong congruency between these assays. Compounds were further evaluated using radiometric enzyme kinases assays using human recombinant STK3, STK4, STK24 (aka MST3) and STK26 (aka MST4) kinase assays (Table 1). Enzymatic IC_50_ values for A1 and B2 were in the low nanomolar range, 41 and 33 nM, respectively.

**Table 1A.** Pyrrolopyridazines (Series A) MST family selectivity and NB correlation with literature compound (XMU-MP-1) for comparison. STK3 IC_50_ and STK3, STK4, 2, 3 and 4 % activity remaining determinations performed in an enzyme assay at Eurofins at the K_m_ of ATP.

| Compound | Alias | STK3 Enzyme IC<sub>50</sub> (nM) | STK3 NB IC<sub>50</sub> (μM) | STK3 % activity remaining (1μM) | STK4 % activity remaining (1μM) | MST3 % activity remaining (1μM) | MST4 % activity remaining (1μM) |
| --- | --- | --- | --- | --- | --- | --- | --- |
| A4 | UNC-BE4-36-1 | 124 | 2.1 |  | 4 | 9 | 51 |
| A3 | UNC-BE4-035 | 7 | 2.0 |  | 2 | 6 | 27 |
| A1 | UNC-BE4-017 | 41 | 7.6 |  | 2 | 4 | 19 |
| A5 | UNC-BE4-040-1 | 391 | >10 |  | 39 | 21 | 74 |
| XMU-MP-1 |  | 635 | 8.4 |  | 14 | 15 | 40 |
| A2 | UNC-BE4-018 | 195 | 9.2 | 15 | 7 | 12 | 54 |
| A6 | UNC-BE4-032 |  | >10 | 44 | 26 | 31 | 74 |
| A7 | UNC-BE4-019 |  | >10 | 62 | 37 | 54 | 52 |
| A8 | UNC-BE4-021 |  | >10 | 70 | 61 | 70 | 30 |
| A9 | UNC-BE4-031 |  | >10 | 82 | 98 | 92 | 106 |
| A10 | UNC-BE4-002 |  | >10 | 86 | 79 | 73 | 78 |
| A11 | UNC-BE4-033 |  | >10 | 89 | 36 | 60 | 71 |
| A12 | UNC-BE4-020 |  | >10 | 92 | 89 | 79 | 86 |
| A13 | UNC-BE4-036 |  | >10 | 138 | 84 | 85 | 82 |

NB = NanoBRET

**Table 1B.** Pyrrolopyrimidine (Series B) MST kinase family selectivity and NB correlation with literature compound (XMU-MP-1) for comparison. STK3 IC_50_ and STK3, STK4, 2, 3 and 4 % activity remaining determinations performed in an enzyme assay at Eurofins at the K_m_ of ATP.

| Compound | Alias | STK3<br>IC<sub>50</sub><br>(nM) | STK3<br>NB IC<sub>50</sub><br>(μM) | STK3 %<br>activity<br>remaining<br>(1μM) | STK4 %<br>activity<br>remaining<br>(1μM) | MST3 %<br>activity<br>remaining<br>(1μM) | MST4 %<br>activity<br>remaining<br>(1μM) |
| --- | --- | --- | --- | --- | --- | --- | --- |
| B6 | UNC-SOB-5-47 | 28 | 0.7 |  | 2 | 24 | 55 |
| B5 | UNC-SOB-5-33 | 57 | 1.6 | 6 | 11 | 14 | 34 |
| B2 | UNC-SOB-5-19 | 33 | 2.7 | 23 | 6 | 36 | 60 |
| B4 | UNC-SOB-5-32 | 142 | 5.0 | 2 | 10 | 62 | 85 |
| B7 | UNC-SOB-5-34 | 53 | 2.3 | 9 | 6 | 33 | 42 |
| B1 | UNC-SOB-5-16 | 113 | 4.3 | 17 | 6 | 29 | 27 |
| B8 | UNC-SOB-5-54 | 283 | 5.7 | 30 | 14 | 54 | 32 |
| B9 | UNC-SOB-5-48 | 80 | 5.0 | 19 | 9 | 51 | 68 |
| B10 | UNC-SOB-5-46 |  | 8.6 | 47 | 32 | 56 | 62 |
| XMU-MP-1 |  | 635 | 8.4 |  | 14 | 15 | 40 |
| B11 | UNC-SOB-5-31 | 299 | >10 | 6 | 12 | 65 | 99 |
| B12 | UNC-SOB-5-18 |  | >10 | 39 | 35 | 74 | 72 |
| B13 | UNC-SOB-5-51 |  | >10 | 49 | 36 | 69 | 81 |
| B14 | UNC-SOB-5-53 |  | >10 | 63 | 44 | 75 | 54 |
| B15 | UNC-SOB-5-49 |  | >10 | 74 | 44 | 80 | 87 |
| B16 | UNC-SOB-5-50 |  | >10 | 76 | 61 | 81 | 55 |
| B17 | UNC-SOB-5-52 |  | >10 | 78 | 59 | 77 | 72 |
| B18 | UNC-SOB-5-29 |  | >10 | 78 | 81 | 78 | 84 |
| B19 | UNC-SOB-5-45 |  | >10 | 104 | 89 | 85 | 85 |
| B3 | UNC-SOB-5-17 |  | >10 | 113 | 61 | 79 | 82 |

NB = NanoBRET

Quantitative densitometry analysis of western blots of p-MOB1 in HMVP2 cells treated with 1 μM for 30 min shows approximately 75% reduction of p-MOB1 in cell lysates treated with A1 and B2 (Fig. 4D). Compounds in series A are derived from JAK inhibitors and accordingly we observed a modest, but significant reduction of p-STAT3 Tyr705 in A1 treated cells only. Compounds in series B are derived from LRRK2 inhibitors; however, phosphorylation of LRRK2 at autophosphorylation site S1292 was not detected in PC cells (24). We tested A1 and B2 against HMVP2 tumor spheroid growth, which have a high level of STK3 expression (Fig. 2A). As shown in Fig. 4E-F, both compounds inhibited HVMP2 spheroid growth by over 50% at 10 μM doses.

### STK3 chemical tools have a protective effect in H9C2 cardiomyocytes

To determine if our investigational compounds are specific to STK3 or have a general toxic effect we utilized the rat H9C2 cardiomyocyte cell line, which are protected from ROS induced apoptosis by activation of YAP including with XMU-MP-1 (25). As shown in Fig. 4G, treatment with increasing doses of A1 and B2 for 72hr had modest growth stimulatory effects on H9C2, nonetheless divergent from effects observed in PC or BC cells. To determine if inhibition of STK3 has a cardio-protective effects we treated cells with a lethal dose of doxorubicin (500 nM) alone or in combination with A1 and B2 (1 μM). Real-time quantitation of cell death showed a steady induction of H9C2 cell death with doxorubicin, which was partially, but significantly, blunted by co-treatment with STK3 inhibitors A1 and B2. Similar chemo-protective effects were observed with the taxane docetaxel (Fig. 4I). These data are consistent with inhibition of canonical Hippo Tumor suppressor function that leads to activation of YAP/TAZ in cardiomyocytes.

### Kinome Selectivity Screening

A1 and B2 were screened for selectivity against a panel of 468 kinases at a 1 μM dose. Both compounds showed a high degree of selectivity (Fig. 4J). A1 inhibited 10 kinases other than STK3 by greater than 90% with an S_10_ (1µM) of 0.027. B2 inhibited only 8 other kinases by >90% of control with an S_10_ (1µM) of 0.022. Kinases with >90% inhibition for A1 and B2 are depicted in Table S1.

### Pyrrolopyrimidine (Series B) analogues have increased potency

Given our encouraging previous results, we proceeded to synthesize and screen additional analogues from both BE4 and SOB-5 scaffolds. Compounds were tested using recombinant protein radiometric assays and in cell STK3 NanoBRET target engagement assays (Table 1 and Fig. S1). From these two assays, compounds with a NanoBRET IC_50_ <2.5 μM and enzymatic IC_50_ < 200 nM were screened for *in vitro* efficacy. Amongst these, we tested B1, B4, B5, B6, A3 and A4 on HMVP2 spheroids at a 5 μM dose (Chemical Structures in Supplemental Data). Over the course of a 9 day HMVP2 spheroid assay, B5 and B6 showed the highest efficacy (Fig. 5A). Western blot analysis of HMVP2 lysates treated with B5 and B6 (1 μM) confirm inhibition of STK3 activity as measured by loss of p-MOB1 T35 to a level of approximately 75% inhibition for both compounds (Fig. 5C-D). We also screened a number of these compounds against off targets identified in our kinome selectivity screen (Fig 4J, Table S1). Consistent with recombinant JAK3 enzyme assays (Table S2), B6, but not B5 significantly reduced levels of JAK3 phospho-target p-STAT3 Tyr705, albeit a modest reduction relative to STK3 inhibition.

**Figure 5.**
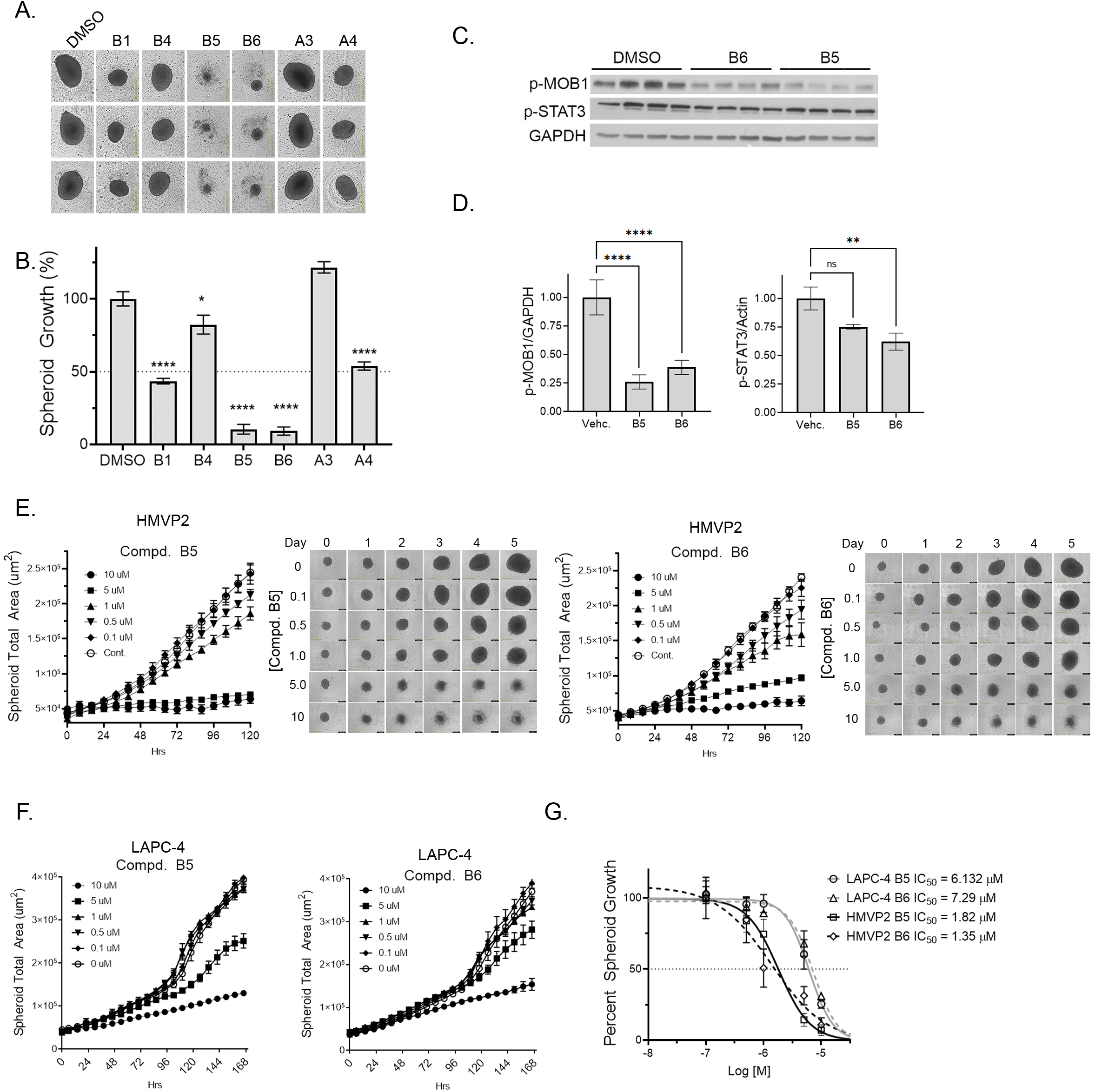
Assessment of lead analogues. A) Representative images of HMVP2 spheroids and B) percent growth relative to vehicle control at endpoint day 9 treated with 5 μM of depicted STK3 inhibitor. C) Western blot analysis of HMVP2 cells treated for 30 min with B5 and B6 at 1uM and D) densitometric analysis normalized to GAPDH. E) HMVP2 spheroid area treated with denoted inhibitor and representative images at day 9. F) LAPC-4 spheroid area treated as denoted at day 9. G) Curve fit of HMVP2 and LAPC-4 spheroid growth assays with B5 and B6 for IC_50_ calculation.

We next conducted dose response curves using STK3 high expressing HMVP2 (Fig. 5E) and STK3 low expressing LAPC-4 (Fig. 5F) spheroid models. Non-linear curve fit to determine IC_50_ values of B5 and B6, which showed lower IC_50_ values in HMVP2 cells compared to LAPC-4 (Fig. 5G).

### STK3 inhibition and loss inhibits PC cell invasion

Next, we tested the effects of STK3 inhibition on in 3D matrigel invasion of HMVP2 cells. In this model, we first tested XMU-MP-1 and our two leads A1 and B2. Compared to vehicle control, HMPV2 spheroids treated with each of the three compounds had statistically significant reduced overall spheroid size and invasion fronts (Fig. S2). We next tested B6, which showed increased STK3 target engagement and efficacy in HMVP2 3D invasion model (Fig. 6A). B6 dramatically reduced overall 3D matrigel invading spheroid area in a dose dependent manner (Fig. 6B). In addition, the invading cell front (blue mask) was universally inhibited at all three doses tested (Fig. 6C). Similarly, genetic depletion of STK3 in HMVP2 cells reduced invasion rates compared to non-targeting control (Fig. 6C-E). Albeit, genetic depletion of STK3 was not as potent as inhibition with B6, likely due to modest STK3 shRNA knockdown in HMVP2 cells. Together, our chemical and genetic data shows that STK3 plays a role in cell invasion.

**Figure 6.**
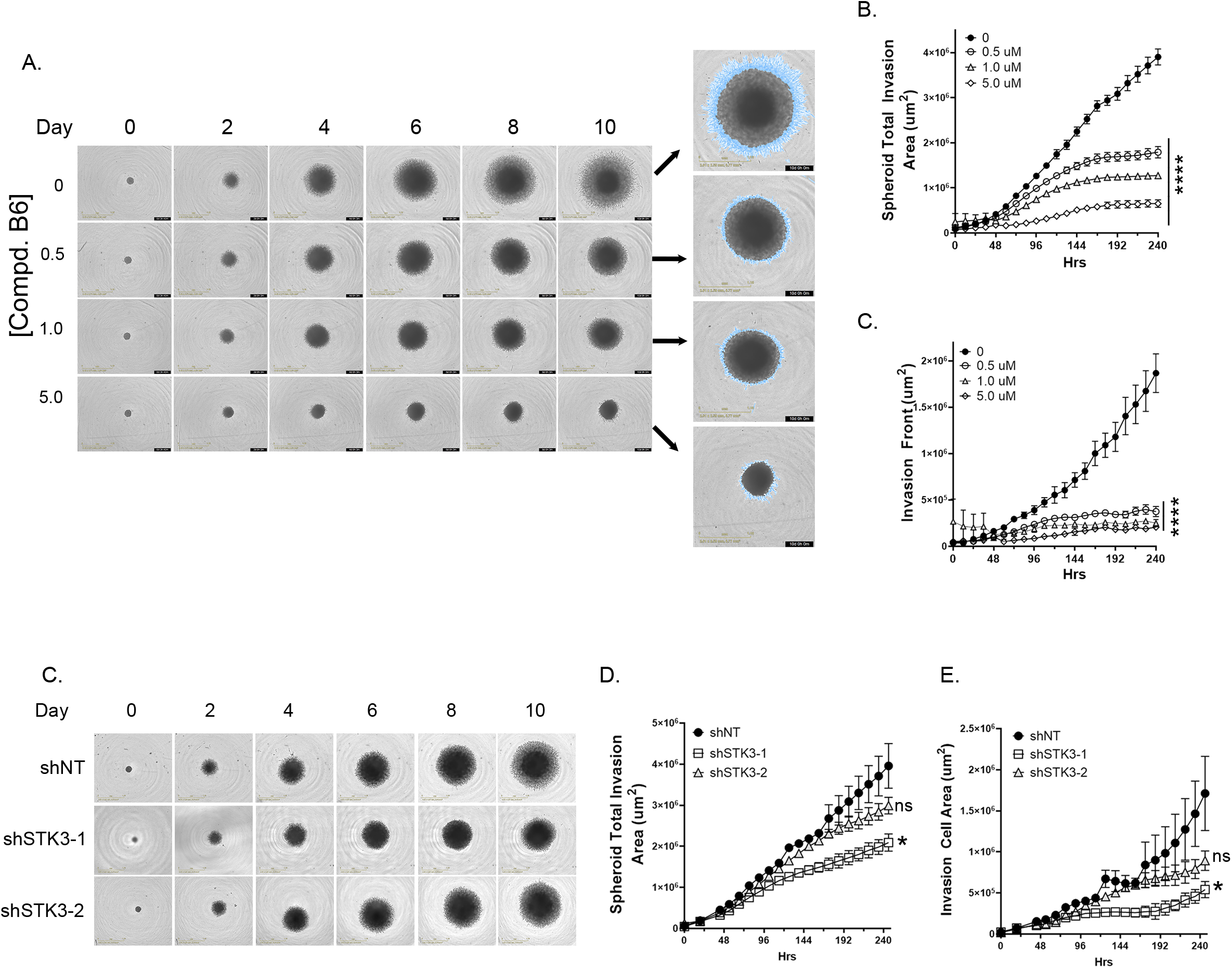
Pharmacological inhibition and genetic depletion of STK3 slows PC cell invasion. A) Representative images of 3D invasion of HMVP2 cells treated as denoted and imaged over time. B) 554 Graphical representation of real-time 3D HMVP2 total spheroid invasion area kinetics and C) cell 555 invasion front (blue mask). C) Representative images of 3D invasion of HMVP2 with denoted shRNAs 556 treated. 2way RM ANOVA *p=0.01, ****p<0.0001.

## Discussion

Our investigation originated from the simple observation that STK3, a known tumor suppressor, is amplified in a subset of cancer types including PC. For this reason, we sought to determine if STK3 plays a non-canonical pro-tumorigenic role in PC. Although this notion contradicts the widely accepted Hippo Tumor suppressor signaling dogma, the combined value of STK3 as a druggable target and the need for added molecular targets to combat mCRPC motivated us to investigate this likelihood. Our study provides genetic and pharmacological evidence that in PC, STK3 is essential for PC cell proliferation and *in vivo* tumor growth.

There is ample evidence that STK3 acts as tumor suppressors in various tumor types other than PC, however data from the literature does support the possibility that STK3 may have a pro-tumorigenic role in a cancer type-dependent manner. For example, Hippo Tumor suppressor kinases LATS1 and LATS2, the immediate downstream phospho-targets of STK3/4, were also shown to be essential for tumorigenesis in a mouse colon cancer model (19). Likewise, a subset of acute myeloid leukemias were found to be dependent on STK3 signaling *in vitro* (18). Lastly, in a retrospective PC study, STK3 expression positively correlated with higher Gleason grade and predicted biochemical recurrence in Taiwanese PC patients (26). Our study expands on the idea that STK3 has a non-canonical role in a cancer type-dependent manner.

Our pharmacological and genetic knockdown studies show that inhibiting STK3 slows PC cell proliferation (*in vitro and in vivo*) and matrigel invasion. The small molecule inhibitors had effects across multiple cell lines with the greatest phenotypic consequences in cell lines with the highest expression of STK3, implicating STK3 as the driver of the response. The impact of the loss of STK3 on invasion is particularly noteworthy as PC metastasis is the main cause of death in patients. A target that is specific to this phenotype in metastatic PC or BC, but not growth inhibit or have cytotoxic effects in normal cells could have pronounced translational impacts. Data from STK3 depleted PC cells and HMVP2 tumors suggest a cell cycle arrest phenotype as shown by accumulation of p27 *in vitro* and loss of p-CDK1/accumulation of cyclin B1 in HMVP2 tumors. This is consistent with STK3 dependent leukemias that suggest STK3 regulates CDK1 to promote cell proliferation (18). In HMVP2 STK3 depleted tumors we did not observe increased cell death as was observed by hyperactive YAP/TAZ due to loss of Hippo kinase (LATS1/2) signaling in a colon cancer model (19). This suggest that in PC cells STK3 pro-tumorigenic role differs from the non-canonical role of LATS1/2 in colon cancer.

Lastly, we found that STK3 gene expression correlates with AR response gene signature in mCRPC. Androgen deprivation (CSS) *in vitro* induced STK3 expression and activity. Importantly loss of STK3 in androgen deprived conditions further slowed cell proliferation. Together this data suggests that STK3 may promote PC progression from hormone sensitive to castrate resistance. Overall, our data are consistent with correlative analysis showing STK3 is frequently amplified in PCs and correlates with worse outcomes in advanced mCRPC.

Our study also presents a new set of small molecule tools to dissect the role and functions of STK3/4. In a screening assay of over 450 kinases, A1 and B2 proved to have a narrow spectrum of activity that is improved compared to XMU-MP-1. While XMU-MP-1 is a potent STK3 inhibitor, it also inhibits 23 other kinases by 90% or more, resulting in a selectivity score S_10_ (1 µM) of 0.05 (17). In comparison, A1 (S_10_ (1 µM) = 0.03) and B2 (S_10_ (1 µM) = 0.02), inhibited 11 and 8 other kinases, respectively, by 90% or more. Our medicinal chemistry campaign yielded compounds with improved efficacy (B5 and B6) in the low micromolar range on 3D tumor spheroids and in the sub-micromolar range on 3D matrigel invasion. Further optimization and *in vivo* evaluation of our compounds is needed, yet at this point these compounds may serve as tools to further investigate STK3 function in different contexts.

To determine potential off target or general toxicity effects of our STK3 kinase inhibitors we tested H9C2 cardiomyocytes. Contrary to our observations in PC cells, STK3 inhibitors did not slow H9C2 cell proliferation. Since STK3 inhibition and activation of YAP have been shown to reduce the effect of apoptotic stimuli in H9C2 cells, we tested the effects of STK3 inhibitors in combination with the commonly used chemotherapeutics doxorubicin and docetaxel. Consistent with a canonical role for STK3, we found chemical inhibition of STK3 with our investigational compounds blunted chemotherapy induced cardiomyocyte cell death. While these are initial observations, it is understood that an STK3 kinase targeted therapy with combined anti-tumor and cardiac chemo-protective effects would hold high clinical significance.

Overall, our study illustrates that STK3 does not always act as a tumor suppressor and that in fact, for a subset of PCs, STK3 provides a pro-tumorigenic growth advantage. Loss of STK3 slowed PC growth and limited invasion potential. We developed potent and specific inhibitors to expand the tools available for probing STK3 and the Hippo/YAP pathway. Importantly, these inhibitors did not slow proliferation in cardiomyocytes and instead provided protection from apoptosis, which illustrates the canonical function of STK3. While deeper mechanistic studies are necessary, our studies support a non-canonical and actionable role for STK3 in PC growth and progression.

