## Supplemental Data and Material/Methods for "Non-canonical role of Hippo tumor suppressor serine/threonine kinase 3 STK3 in prostate cancer"

**Supplemental Data Schirmer et. al. 2021**

Table S1. Kinases with <10% activity remaining as profiled in KINOME*scan* panel at 1 μM

| **Compound A1** | | | **Compound B2** | |
|---|---|---|---|---|
| Kinase | PoC | | Kinase | PoC |
| CSNK2A1 | 0 | | LRRK2 | 0 |
| MAP3K15 | 0 | | LRRK2 (G2019S | 0 |
| STK3 (MST2)* | 0 | | STK3 (MST2) | 0 |
| RIOK2 | 0 | | MAP4K3 | 3.4 |
| JAK2 | 0.4 | | BIKE | 3.8 |
| JAK3 | 0.7 | | PIP5K1A | 4.2 |
| NUAK2 | 1.2 | | NIK | 4.4 |
| BMPR1B | 1.8 | | NUAK2 | 7.5 |
| LRRK2 | 2.1 | | YSK4 | 7.6 |
| NUAK1 | 4.1 | | GRK4 | 8.1 |
| TYK2 | 6.9 | | | |
| LRRK2 (G2019S) | 7.5 | | | |

PoC: Percent of Control
Compounds screened at 1 μM

*Different companies screen kinases under different aliases.

| **Kinase** | **Alias** |
|---|---|
| STK3 | MST2 |
| STK4 | MST1 |
| STK24 | MST3 |
| MST4 | |

Table S3. Off targets of pyrrolopyrimidine STK3 inhibitors

| | B6 | B2 | B7 | B5 | B4 |
|---|---|---|---|---|---|
| **STK3 Enzyme $IC_{50}$** | 28 nM | 33 nM | 53 nM | 57 nM | 142 nM |
| **STK3 NB $IC_{50}$** | 0.7 µM | 2.7 µM | 2.3 µM | 1.6 µM | 5.0 µM |

| Kinase | B6 | B2 | B7 | B5 | B4 | [ATP] µM |
|---|---|---|---|---|---|---|
| LRRK2 | 5 | 0 | 2 | 4 | 5 | **70** |
| MAP4K3 | 2 | 0 | 12 | 11 | 15 | **45** |
| GCK (MAP4K2) | 5 | 2 | 18 | 17 | 45 | **200** |
| BIKE | 11 | 15 | 9 | 7 | 24 | **10** |
| NUAK2 | 6 | 26 | 17 | 14 | 35 | **15** |
| RSK4 | 13 | 40 | 20 | 26 | 62 | **45** |
| TAO1 | 35 | 57 | 46 | 34 | 68 | **10** |
| JAK3 | 5 | 73 | 53 | 46 | 63 | **10** |
| MEKK3 | 39 | 74 | 83 | 91 | 102 | **10** |
| MEK1 | 56 | 76 | 55 | 63 | 112 | **10** |
| MSSK1 | 71 | 94 | 84 | 83 | 86 | **15** |
| PIP5K1a | 104 | 101 | 103 | 100 | 101 | **200** |
| BTK | 106 | 104 | 83 | 99 | 91 | **200** |
| PRP4 | 107 | 112 | 105 | 92 | 119 | **15** |

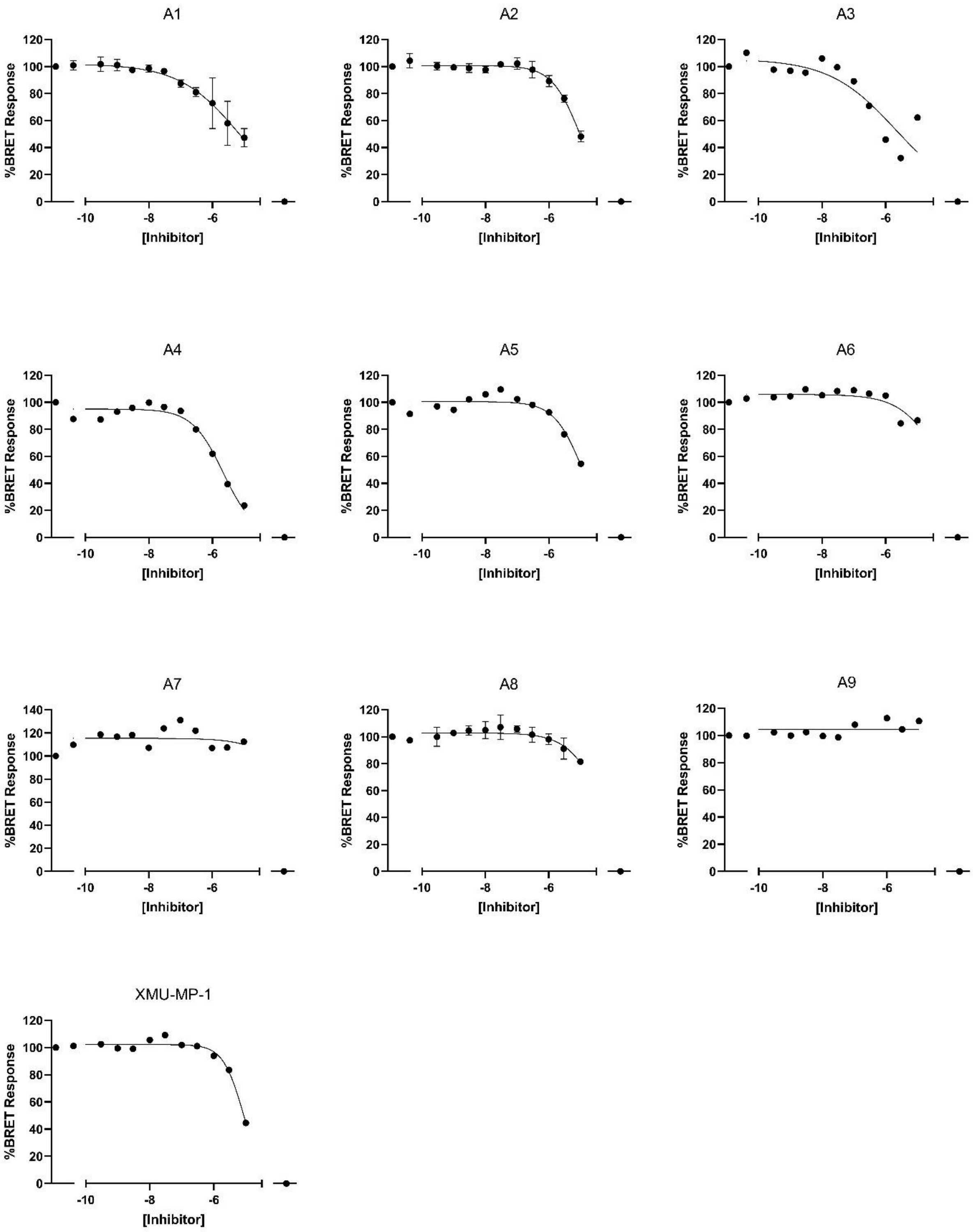


**Figure S1. Series A normalized STK3 NanoBRET $IC_{50}$ graphs. (Graphpad Prism)**

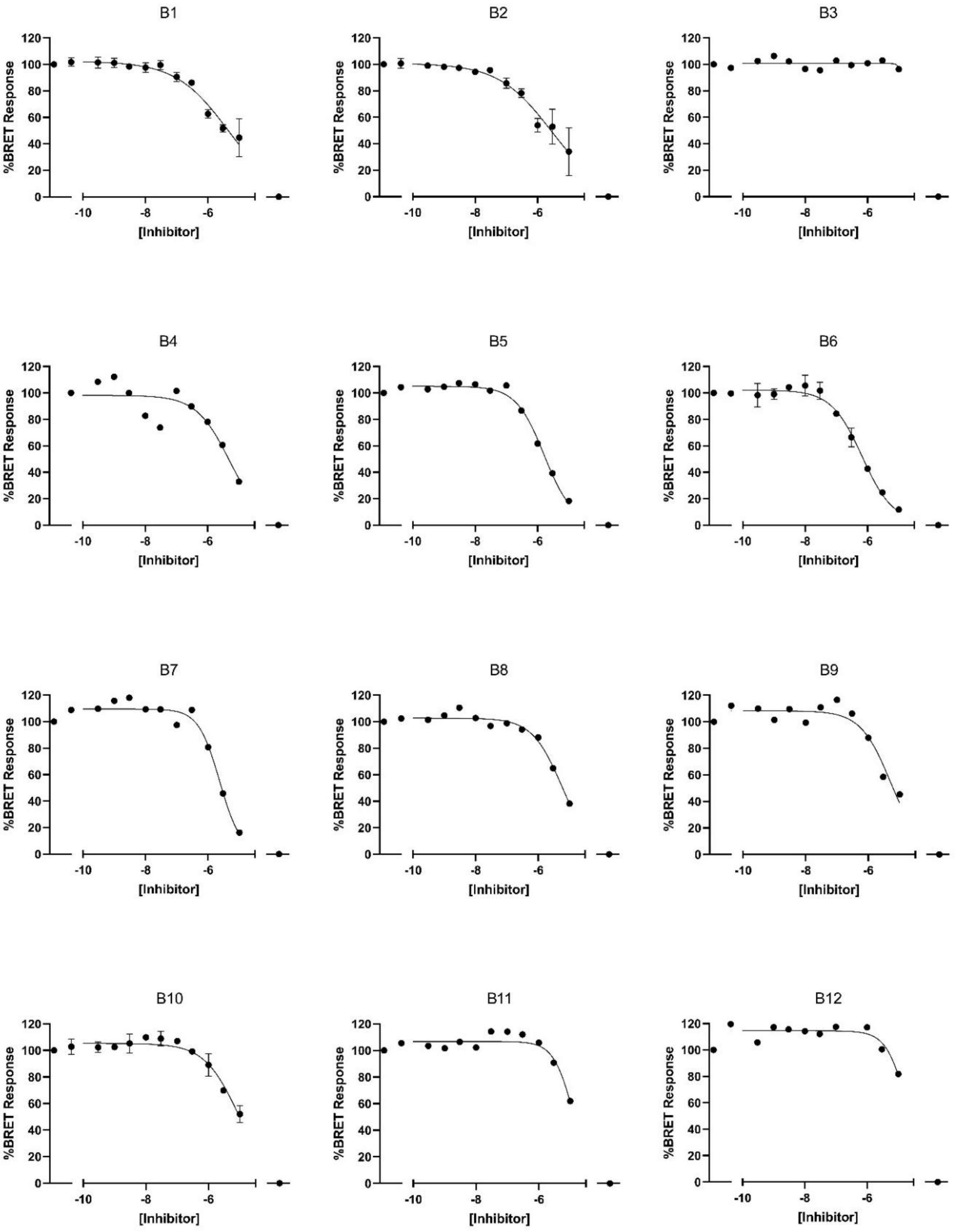
B1
B2
B3
B4
B5
B6
B7
B8
B9
B10
B11
B12
%BRET Response
[Inhibitor]
0
20
40
60
80
100
120
-10
-8
-6

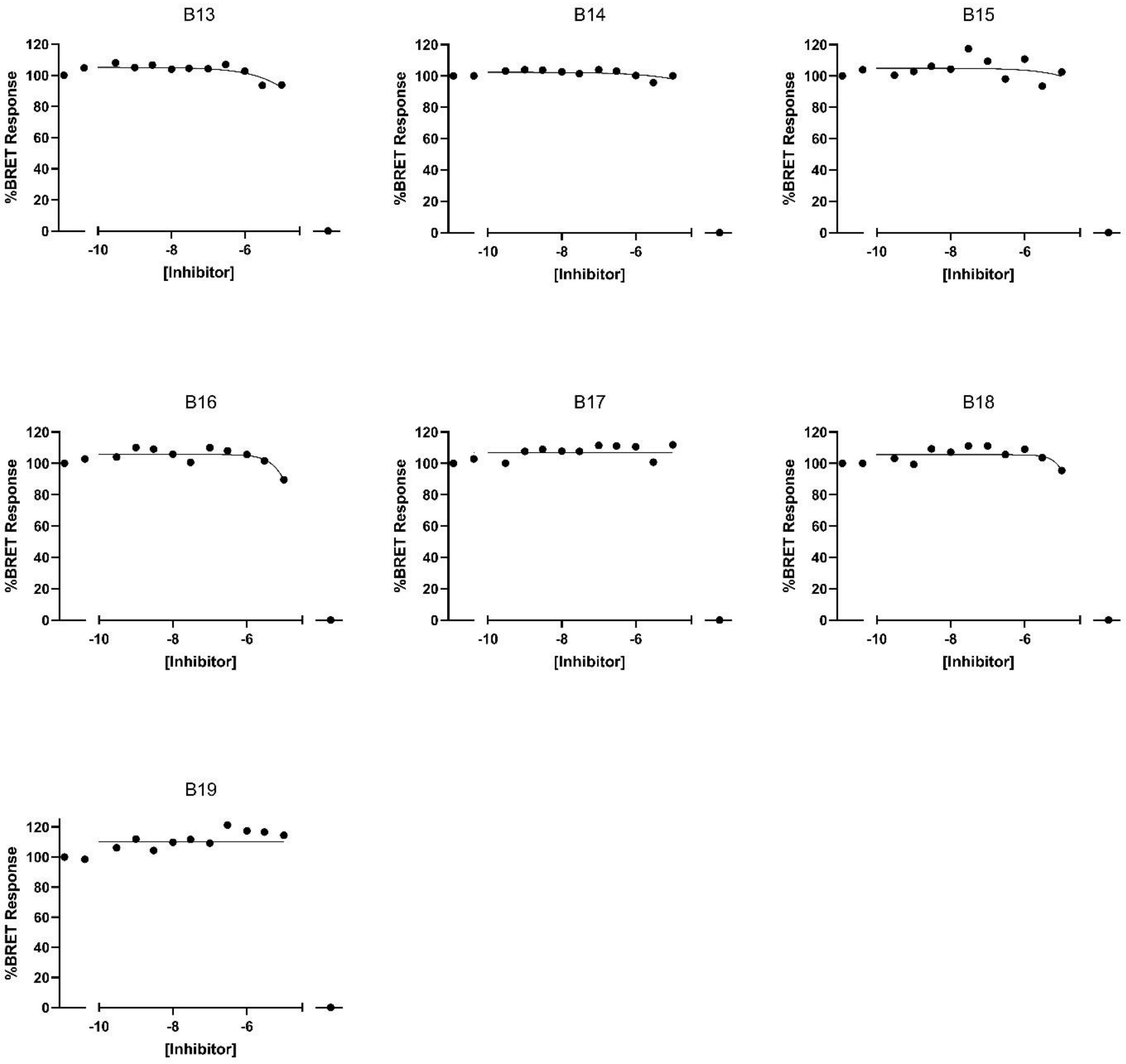


Figure S1. Series B normalized STK3 NanoBRET $IC_{50}$ graphs. (Graphpad Prism)

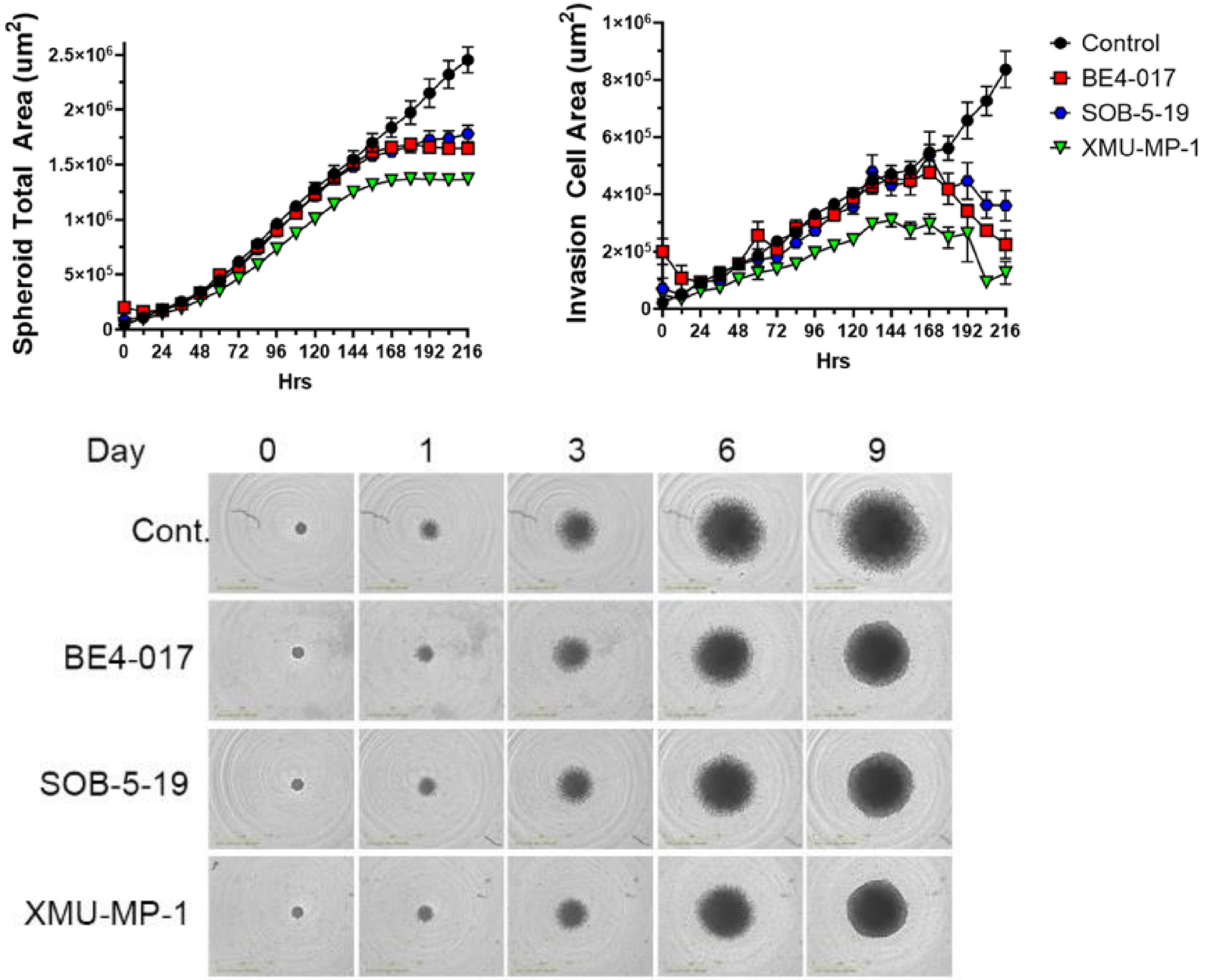


Figure S2. HMVP2 3D spheroid invasion treated with STK3 inhibitors.

### Supplemental Materials and Methods: Synthetic Routes and Compound Characterization

*General chemistry information*: All reagents and solvents, unless specifically stated, were used as obtained from their commercial sources without further purification. Solvents were degassed with nitrogen for cross-coupling reactions. Air and moisture sensitive reactions were performed under an inert atmosphere using nitrogen in a previously oven-dried or flame-dried reaction flask, and addition of reagents were done using a syringe. All microwave (µW) reactions were carried out in a Biotage Initiator EXP US 400W microwave synthesizer. Thin layer chromatography (TLC) analyses were performed using 200 µm pre-coated sorbtech fluorescent TLC plates and spots were visualized using UV light. High resolution mass spectrometry samples were analyzed with a ThermoFisher Q Exactive HF-X (ThermoFisher, Bremen, Germany) mass spectrometer coupled with a Waters Acquity H-class liquid chromatograph system. All HRMS were obtained via electrospray ionization (ESI). Column chromatography was undertaken with a Biotage Isolera One or Prime instrument. Nuclear magnetic resonance (NMR) spectrometry was run on a varian Inova 400 MHz or Bruker Avance III 700 MHz spectrometer equipped with a TCI H-C/N-D 5 mm cryoprobe and data was processed using the MestReNova processor. Chemical shifts are reported in ppm with residual solvent peaks referenced as internal standard.

#### A-series: amido-pyrrolopyrazines

The compounds in the pyrrolopyrazine series (A-series) were synthesized as described in 3-Amido Pyrrolopyrazine JAK Kinase Inhibitors: Development of a JAK3 vs. JAK1 Selective Inhibitor and Evaluation in Cellular and In-Vivo Models" J Med Chem (2013) **56**, p.345. PMID: 23214979.

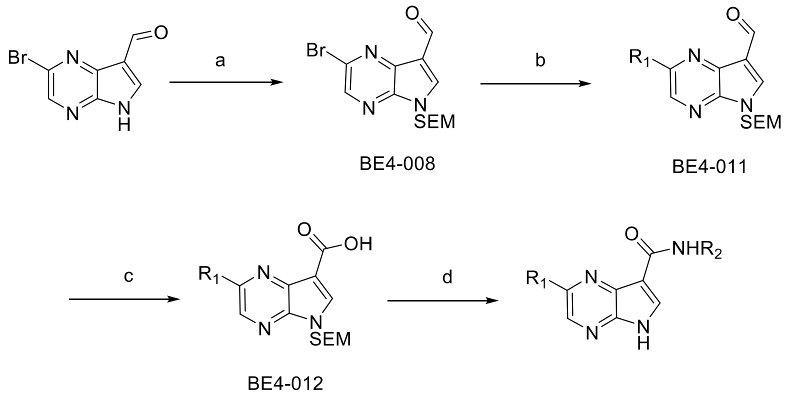


**Reagents and conditions**: a) NaH, dimethylformamide, 0-25 $^{o}$C, 1.5 h; b) $R_1$ [Cyclopropylboronic acid or phenylboronic acid], $K_3PO_4$, $Pd(OAc)_2$, $P(Cy)_3$, Toluene/water, 100 $^{o}$C, 18 h; c) $KH_2PO_4$, $NaClO_2$, $H_3NSO_4$, 1,4-dioxane/water, 2 h; d) i. $R_2$-$NH_2$, HATU, DIPEA, acetonitrile, 16 h, ii. TFA, dichloromethane, 2 h., iii. Sodium acetate, ethanol, 20 h.

**2-bromo-5-((2-(trimethylsilyl)ethoxy)methyl)-5H-pyrrolo[2,3-*b*]pyrazine-7-carbaldehyde** (**BE4-008**). To a solution of 2-bromo-5H-pyrrolo[2,3-*b*]pyrazine-7-

carbaldehyde (4.7 g, 21 mmol) in DMF (50 mL) at 0 °C was added 60% sodium hydride (1.5 g, 62 mmol) in mineral oil. The reaction mixture was stirred at 25 °C for 30 min and then cooled to 0 °C. The mixture was then treated with (2-(chloromethoxy)ethyl)trimethylsilane (4.4 mL, 25 mmol). The reaction was allowed to warm to 25 °C, stirred for 1 h, and then quenched with water and extracted three (3) times with ethyl acetate. The combined organics were sequentially washed with three portions of water, and a saturated sodium chloride solution and then dried over sodium sulfate, filtered and concentrated. The residue was purified by silica gel chromatography (20-30% ethyl acetate/hexanes) to afford 2-bromo-5-((2-(trimethylsilyl)ethoxy)methyl)-5H-pyrrolo[2,3-*b*]pyrazine-7-carbaldehyde (6.3 g, 18 mmol, 85 %) as a yellow solid. This material was used in the next step without further characterization. LCMS $[M+H]^+$: m/z 355.2/357.2

**2-cyclopropyl-5-((2-(trimethylsilyl)ethoxy)methyl)-5H-pyrrolo[2,3-*b*]pyrazine-7-carbaldehyde** (**BE4-011**). A mixture of 2-bromo-5-((2-(trimethylsilyl)ethoxy)methyl)-5H-pyrrolo[2,3-*b*]pyrazine-7-carbaldehyde (1.2 g, 3.4 mmol), cyclopropylboronic acid (0.43 g, 5.1 mmol), tricyclohexylphosphine (94 mg, 0.34 mmol) , palladium(II) acetate (38 mg, 0.17 mmol) , potassium phosphate (2.3 g, 11 mmol) , toluene (16 mL), Water (2.0 mL) was flushed with nitrogen for 5 mins. The reaction mixture was stirred at 100 °C for 18 h and then allowed to cool and filtered through a pad of Celite 545, rinsing with ethyl acetate. The filtrate was concentrated under reduced pressure, and the residue was purified by silica gel chromatography (10% ethyl acetate/hexane) to afford 2-cyclopropyl-5-((2-(trimethylsilyl)ethoxy)methyl)-5H-pyrrolo[2,3-*b*]pyrazine-7-carbaldehyde (0.95 g, 89%) as a yellow powder. $^1H$ NMR (400 MHz, DMSO-d6) δ 10.07 (s, 1H), 8.78 (s, 1H), 8.43 (s, 1H), 5.68 (s, 2H), 3.59 – 3.53 (m, 2H), 2.34 (ddd, J = 7.9, 4.9, 2.8 Hz, 1H), 1.05 (tt, J = 7.7, 2.6 Hz, 4H), 0.86 – 0.80 (m, 2H), -0.10 (s, 8H). LCMS $[M+H]^+$: m/z 317.5

**2-phenyl-5-((2-(trimethylsilyl)ethoxy)methyl)-5H-pyrrolo[2,3-*b*]pyrazine-7-carbaldehyde** (**BE4-011-PHENYL**). Using conditions similar to the one outlined above, 2-phenyl-5-((2-(trimethylsilyl)ethoxy)methyl)-5H-pyrrolo[2,3-b]pyrazine-7-carbaldehyde was obtained as a pale solid; 92%. LCMS $[M+H]^+$: m/z 353.5.

**2-cyclopropyl-5-((2-(trimethylsilyl)ethoxy)methyl)-5H-pyrrolo[2,3-b]pyrazine-7-carboxylic acid** (**BE4-012**). To a mixture of 2-cyclopropyl-5-((2-(trimethylsilyl)ethoxy)methyl)-5H-pyrrolo[2,3-*b*]pyrazine-7-carbaldehyde (800 mg, 2.52 mmol) in 1,4-Dioxane (25 mL) and Water (5.0 mL) at 0 °C was added sulfamic acid (1.47 g, 15.1 mmol) and then dropwise a solution of 80% sodium chlorite (251 mg, 2.77 mmol) and potassium dihydrogen phosphate (4.12 g, 30.2 mmol) in 18 mL of water. The reaction was allowed to warm to rt and stirred for 2 h and partitioned between water and ethyl acetate. The organic phase was washed with a saturated aqueous sodium chloride, dried over sodium sulfate, filtered, and concentrated under reduced pressure. The residue was triturated with hexanes to obtain 2-cyclopropyl-5-((2-(trimethylsilyl)ethoxy)methyl)-5H-pyrrolo[2,3-*b*]pyrazine-7-carboxylic acid (0.70 g, 83 %) as a light yellow powder. LCMS $[M+H]^+$: m/z 333.5

| Compound | | $R_1$ | $R_2$ |
|---|---|---|---|
| **A1** | **UNC-BE4-017** | 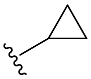 | 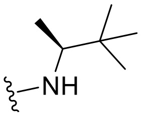 |
| **A2** | **UNC-BE4-018** | 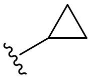 | 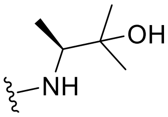 |
| **A3** | **UNC-BE4-035-1** | 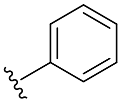 | 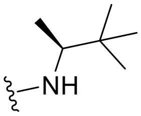 |
| **A4** | **UNC-BE4-036-1** | 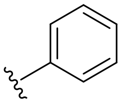 | 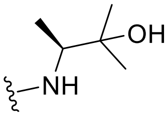 |
| **A5** | **UNC-BE4-040** | 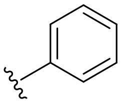 | 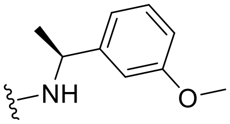 |
| **A6** | **UNC-BE4-032** | 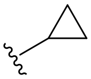 | 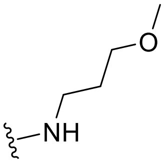 |
| **A9** | **UNC-BE4-031** | 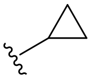 | 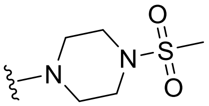 |
| **A11** | **UNC-BE4-033** | 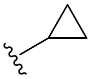 | 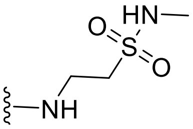 |
| **A13** | **UNC-BE4-036** | 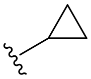 | 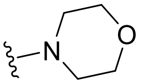 |

$^1$H NMR Characterization data of key A-series compounds:

**A1 (UNC-BE4-017).** (S)-2-cyclopropyl-N-(3,3-dimethylbutan-2-yl)-5H-pyrrolo[2,3-*b*]pyrazine-7-carboxamide (34 mg, 78 %). $^1$H NMR (400 MHz, DMSO-d6) δ 12.54 (s,

1H), 8.40 (s, 1H), 8.25 (s, 1H), 8.14 (d, J = 9.4 Hz, 1H), 3.92 (dq, J = 9.4, 6.7 Hz, 1H), 2.34 (tt, J = 8.1, 4.8 Hz, 1H), 1.11 (d, J = 6.8 Hz, 4H), 1.09 – 0.91 (m, 12H).

**A2 (UNC-BE4-018).** $^1$H NMR (400 MHz, DMSO-d6) δ 12.49 (s, 1H), 8.39 (d, J = 11.0 Hz, 2H), 8.23 (s, 1H), 4.58 (s, 1H), 3.92 (dq, J = 8.8, 6.6 Hz, 1H), 2.31 (tt, J = 7.9, 5.1 Hz, 1H), 1.20 – 1.12 (m, 9H), 1.11 – 1.00 (m, 4H).

**A3 (UNC-BE4-035-1).** $^1$H NMR (400 MHz, DMSO-$d_6$) δ 12.78 (s, 1H), 8.99 (s, 1H), 8.40 (d, *J* = 8.0 Hz, 2H), 8.17 – 8.10 (m, 2H), 7.60 – 7.52 (m, 2H), 7.52 – 7.44 (m, 1H), 3.98 (dq, *J* = 9.2, 6.7 Hz, 1H), 1.16 (d, *J* = 6.8 Hz, 3H), 1.00 (s, 9H).

**A4 (UNC-BE4-036-1).** $^1$H NMR (400 MHz, DMSO-$d_6$) δ 12.73 (s, 1H), 9.02 (s, 1H), 8.71 (d, *J* = 8.7 Hz, 1H), 8.38 (s, 1H), 8.31 – 8.27 (m, 2H), 7.57 – 7.44 (m, 3H), 4.73 (s, 1H), 4.01 – 3.94 (m, 1H), 1.27 – 1.14 (m, 9H).

**A5 (UNC-BE4-040).** $^1$H NMR (400 MHz, DMSO-$d_6$) δ 12.79 (s, 1H), 9.02 (s, 1H), 8.78 (d, *J* = 7.5 Hz, 1H), 8.43 (s, 1H), 8.13 – 8.08 (m, 2H), 7.57 – 7.42 (m, 3H), 7.32 (t, *J* = 7.9 Hz, 1H), 7.11 – 7.04 (m, 2H), 6.89 (ddd, *J* = 8.2, 2.6, 0.9 Hz, 1H), 5.18 (t, *J* = 7.0 Hz, 1H), 3.73 (s, 3H), 1.58 (d, *J* = 6.8 Hz, 3H).

**A6 (UNC-BE4-032).** $^1$H NMR (400 MHz, DMSO-$d_6$) δ 12.54 (s, 1H), 8.37 (s, 1H), 8.26 (s, 1H), 8.16 (t, *J* = 5.7 Hz, 1H), 3.44 (q, *J* = 6.2 Hz, 4H), 3.27 (s, 3H), 2.33 (tt, *J* = 8.0, 4.8 Hz, 1H), 1.78 (p, *J* = 6.5 Hz, 2H), 1.11 – 1.00 (m, 4H).

**A9 (UNC-BE4-031).** $^1$H NMR (400 MHz, DMSO-$d_6$) δ 12.51 (s, 1H), 8.33 (s, 1H), 8.16 (s, 1H), 3.70 (t, *J* = 5.5 Hz, 4H), 3.23 (t, *J* = 5.1 Hz, 4H), 2.92 (s, 3H), 2.30 (tt, *J* = 8.0, 4.9 Hz, 1H), 1.05 – 0.97 (m, 4H).

**A11 (UNC-BE4-033).** $^1$H NMR (400 MHz, DMSO-$d_6$) δ 12.57 (s, 1H), 8.37 (d, *J* = 15.3 Hz, 2H), 8.29 (s, 1H), 7.07 (q, *J* = 4.9 Hz, 1H), 3.76 (q, *J* = 6.2 Hz, 2H), 3.28 (t, *J* = 6.3 Hz, 2H), 2.61 (d, *J* = 4.9 Hz, 3H), 2.30 (tt, *J* = 8.2, 4.8 Hz, 1H), 1.18 – 1.13 (m, 2H), 1.03 (dt, *J* = 8.3, **3.1 Hz, 2H).**

**A13 (UNC-BE4-036).** $^1$H NMR (400 MHz, DMSO-$d_6$) δ 12.49 – 12.44 (m, 1H), 8.31 (s, 1H), 8.13 (s, 1H), 3.67 (t, *J* = 5.5, 3.8 Hz, 4H), 3.57 (t, *J* = 4.7 Hz, 4H), 2.29 (tt, *J* = 8.1, 4.8 Hz, 1H), 1.06 – 0.92 (m, 4H).

**B-series: pyrrolopyrimidines**

The compounds in the pyrrolopyrimidine series (B-series) were synthesized as described in "Discovery and preclinical profiling of 3-[4-(morpholin-4-yl)-7H-pyrrolo[2,3-d]pyrimidin-5-yl]benzonitrile (PF-06447475), a highly potent, selective, brain penetrant, and in vivo active LRRK2 kinase inhibitor" J Med Chem (2015) **58**, p.419. PMID: 25353650.

General scheme:

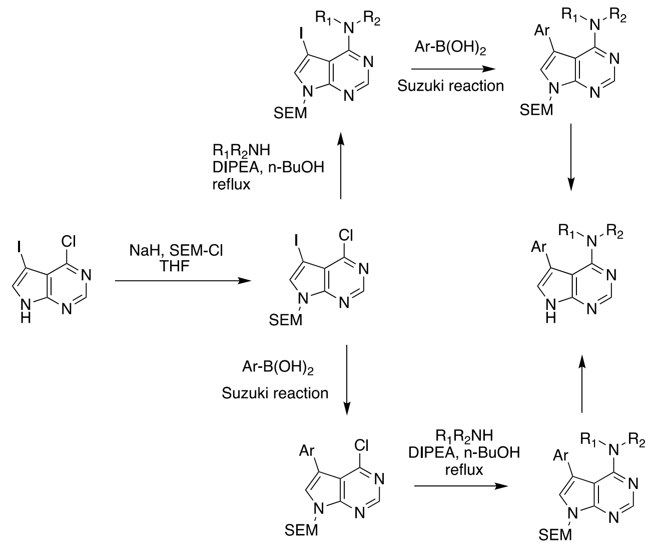


$^1$H NMR Characterization data of key B-series compounds:

**B1 (UNC-SOB-5-16)**

$^1$H NMR (400 MHz, MeOD) δ 8.26 (s, 1H), 7.85 (t, *J* = 1.5 Hz, 1H), 7.79 (dt, *J* = 7.56, 1.53, 1H), 7.65 (dt, *J* = 7.74, 1.44 Hz, 1H), 7.59 (t, *J* = 7.63, H), 7.37 (s, 1H), 2.82 (s, 6H)

**B2 (UNC-SOB-5-19)**

$^1$H NMR (400 MHz, DMSO-*d*6) δ 12.35 (s, 1H) 8.40 (s, 1H), 8.00 (s, 1H), 7.89 (d, *J* = 7.9 Hz, 1H), 7.76 (d, *J* = 7.8 Hz, 1H), 7.70 (s, 1H), 7.67 (t, *J*= 7.8 Hz, 1H), 3.47 (t, *J* = 4.51, 4H), 3.14 (t, *J* = 4.56, 4H).

**B3 (UNC-SOB-5-17)**

$^1$H NMR (400 MHz, $CDCl_3$) δ 8.33 (d, *J* = 13.4 Hz, 1H), 7.54 (d, *J* = 1.6 Hz, 1H), 7.44 (d, *J* = 3.3 Hz, 1H), 7.08 (d, *J* = 10.0 Hz, 1H), 3.97 (app d, *J* = 2.4 Hz, 3H), 2.99 (s, 3H), 2.92 (s, 3H)

**B4 (UNC-SOB-5-32)**

$^1$H NMR (700 MHz, DMSO-*d*6) δ 12.17 (s, 1H), 8.38 (s, 1H), 8.09 (s, 1H), 8.01 (s, 1H), 7.80 (d, *J* = 7.5 Hz, 1H), 7.69 (d, *J* = 7.4 Hz, 1H), 7.57 (s, 1H), 7.52 (t, *J* = 7.4 Hz, 1H), 7.37 (s, 1H), 3.42 (s, 4H), 3.14 (s, 4H)

**B5 (UNC-SOB-5-33)**

$^1$H NMR (400 MHz, DMSO-*d*6) δ 11.99 (s, 1H), 9.40 (s, 1H), 8.32 (s, 1H), 7.32–7.38 (m, 2H), 3.47–3.42 (m, 4H), 3.21–3.14 (m, 4H)

**B6 (UNC-SOB-5-47)**

$^{1}H$ NMR (700 MHz, MeOD) δ 8.31 (s, 1H), 7.27–7.23 (m, 2H), 6.98 (d, *J* = 7.3 Hz, 1H), 6.93 (s, 1H), 6.76 (d, *J* = 7.9 Hz, 1H), 3.54 (s, 4H), 3.29 (s, 4H)
